# The genus *Cortinarius* should not (yet) be split

**DOI:** 10.1101/2024.05.29.596376

**Authors:** Brigida Gallone, Thomas W. Kuyper, Jorinde Nuytinck

**Affiliations:** Naturalis Biodiversity Center, Darwinweg 2, 2333 CR Leiden, the Netherlands; Soil Biology Group, Wageningen University, 6700 AA Wageningen, the Netherlands; Research group Mycology, Department of Biology, Ghent University, K.L. Ledeganckstraat 35, 9000 Ghent, Belgium

**Keywords:** Phylogenomics, Classification, Nomenclature, Phylogenetic conflict

## Abstract

The genus *Cortinarius* (Pers.) Gray (Agaricales, Basidiomycota) is one of the most species-rich fungal genera with thousands of species reported. *Cortinarius* species are important ectomycorrhizal fungi and form associations with many vascular plants globally. Until recently *Cortinarius* was the single genus of the *Cortinariaceae* family, despite several attempts to provide a workable, lower-rank hierarchical structure based on subgenera and sections. The first phylogenomic study for this group elevated the old genus *Cortinarius* to family level and the family was split into ten genera, of which seven were described as new. Here, by careful re-examination of the recently published phylogenomic dataset, we detected extensive gene-tree/species-tree conflicts using both concatenation and multispecies coalescent (MSC) approaches. Our analyses demonstrate that the *Cortinarius* phylogeny remains unresolved and the resulting phylogenomic hypotheses suffer from very short and unsupported branches in the backbone. We can confirm monophyly of only four out of ten suggested new genera, leaving uncertain the relationships between each other and the general branching order. Thorough exploration of the tree space demonstrated that the topology on which *Cortinarius* revised classification relies on does not represent the best phylogenetic hypothesis and should not be used as constrained topology to include additional species.

For this reason, we argue that based on available evidence the genus *Cortinarius* should not (yet) be split.

Moreover, considering that phylogenetic uncertainty translates to taxonomic uncertainty, we advise for careful evaluation of phylogenomic datasets before proposing radical taxonomic and nomenclatural changes.

## Introduction

Phylogenetic inference based on genome-scale data has led to significant improvements in the reconstruction of robust phylogenetic hypotheses for diverse sections of the fungal tree of life, especially at the level of phyla, subphyla, and classes (Shen et al. 2016b, 2020; Grewe et al. 2020; Li et al. 2021; Strassert and Monaghan 2022).

Nevertheless, complex evolutionary histories remain difficult to disentangle despite the availability of increasingly comprehensive datasets, and it has become clear that 1) more data does not necessarily result in less incongruence and conflict in phylogenetic analyses (Philippe et al. 2011; Shen et al. 2016a, 2017; Reddy et al. 2017; Jorna et al. 2021a) and 2) traditional statistical branch support measures (e.g. bootstrap) fail to address the nature of variation in these datasets (Chan et al. 2020; van de Peppel et al. 2021).

Discordant phylogenetic signal across loci in a genome-scale dataset can be caused by technical issues such as biased/poor taxon sampling, gene choice, model selection, and biological processes including rapid diversification, incomplete lineage sorting, gene duplication and introgression (for a review see (Steenwyk et al. 2023)). Therefore, careful evaluation of the discordant signal across loci and methodologies is a critical step in the estimation of the most likely phylogenetic hypothesis.

Robust phylogenies are a fundamental prerequisite for evolutionary analyses (Leavitt et al. 2016; Lofgren et al. 2021; Tian et al. 2021; Sierra-Patev et al. 2023). Phylogenetic uncertainty and instability have important consequences on the study of macroevolutionary patterns, diversification, trait evolution and ecological processes (Misof et al. 2014; Jarvis et al. 2014; Parks et al. 2018; Varga et al. 2019; Leebens-Mack et al. 2019; Ametrano et al. 2019; Kawahara et al. 2019).

Most importantly, the significance of phylogenetic uncertainty becomes evident when it translates to taxonomic uncertainty (Xu 2020; Chafin et al. 2021; Jorna et al. 2021b). In this context uncertainty arises from non-monophyletic grouping, unstable relationships between assumed monophyletic groupings, instability due to the addition of further species or a change in outgroup choice. To some extent uncertainty is due to incomplete or biased sampling(Vellinga et al. 2015; Stengel et al. 2022). However, that is likely only part of the explanation.

In mushroom-forming Basidiomycota, the use of genome-scale data applied to taxonomic classification and revision is still in its infancy but rapidly evolving. Species-rich fungal genera are particularly challenging for taxonomists and ecologists (Kandawatte Wedaralalage et al. 2020; Bhunjun et al. 2022; Phukhamsakda et al. 2022). Major challenges refer to the need to provide a workable hierarchical structure of that taxon, the recognisability of species based on morphological and / or molecular characteristics, the rates of speciation and extinction that resulted in such large species numbers, and the niches that the various species occupy and that allow them to co-exist.

The genus *Cortinarius* (Pers.) Gray (Agaricales, Basidiomycota) is one of the most species-rich fungal genera and for this reason exemplifies this challenge. In the kingdom of Fungi it was ranked as the second-most species-rich genus with 3059 species as listed in Species Fungorum by 2021 (Bhunjun et al. 2022). That number is almost certainly an underestimate. Based on available ITS sequences in public databases it was estimated the existence of 1168 new, still undescribed species and the actual number could be above 5000 species (Bhunjun et al. 2022). Recently, a global soil DNA metabarcoding survey of fungal endemicity ranked *Cortinarius* the sixth largest genus with 14375 operational taxonomic units reported (full-length ITS regions clustered at 98% sequence similarity) and used as proxy for species (Tedersoo et al. 2022).

Species of *Cortinarius* form ectomycorrhizas and their host range varies from extremely specialised (with associations with one plant genus only) to rather generalist with associations with members of a larger number of genera, including coniferous and broad-leaved trees, shrubs and some herbaceous plants. They are globally distributed from the tropics to arctic and alpine regions in both hemispheres, with some species occurring almost globally and others being almost local endemics. However, as far as known, species either occur in the northern or southern hemisphere; cases where species have been reported from both hemispheres refer to cases of human introductions (Peintner et al. 2004; Garnica et al. 2005).

Most members of the genus *Cortinarius* are easily recognisable by the combination of a rusty brown spore print (Matheny et al. 2015), ornamented spores and the presence of a cortina, a cobweb-like structure that connects the pileus with the stipe in young specimens. For that reason, Fries (1851) considered the genus as a natural (in the pre-Darwinian sense of common essence, not common descent) genus.

Because of its large size Fries (1838) already divided *Cortinarius* in six subgroups. These subgroups have sometimes been recognised as genera, although most mycologists preferred to recognise one large genus. However, as these subgroups are fairly easily recognisable, at least in the northern hemisphere, these groups often persisted in identification keys. Integrating species from the southern hemisphere turned out to be challenging, and the problems were resolved by the addition of a few more subgroups rather than through a radical modification of an infrageneric classification (Moser et al. 1975).

The advent of molecular data enabled to revisit the existing *Cortinarius* classification. DNA barcoding confirmed the monophyly of *Cortinarius* and also revealed that the traditionally recognised subgroups were polyphyletic. No attempts were, however, made to propose a new classification based on monophyletic subgroups. The first systematic attempt to arrive at a phylogenetically supported new classification of *Cortinarius* was undertaken by Peintner et al. (2004) (Peintner et al. 2004). They noted that their molecular-sequence data demonstrated that species of *Cortinarius*, although morphologically quite variable, are conserved by comparison in their ribosomal DNA (rDNA) region. Short backbone branches characterized their phylogeny. They confirmed that *Cortinarius* was monophyletic and did not formally propose a new classification. They ended their paper with the question ‘Can the *Cortinarius* phylogeny be resolved?’, providing a cautious and provisional ‘not yet’ answer. A subsequent study with increased taxon sampling (Garnica et al. 2005), especially with southern-hemisphere species, confirmed some of the previous results but also resulted in notable differences. That study did not indicate whether the changed topology was due to their species selection or to the choice of a different outgroup. The study also did not propose a new formal classification. Soop et al. (2019) further enlarged the number of species and used additional genes beyond those of the ribosomal cluster. They formally recognised 79 clades that they described as sections and recovered an additional 20 clades where they refrained from formal description. No further taxonomic hierarchy was proposed. They stated that for a more complete hierarchical framework that uses these sections as building blocks for a complete hierarchical structure a larger number of genes or preferably a phylogenomics approach would be required. That step was recently taken by Liimatainen et al. (2022) (Liimatainen et al. 2022). They reported to have performed phylogenomic analysis for 19 species based on 75 single-copy genes; and a subsequent, constrained analysis for 245 species based on five single-copy genes. Consequently, the genus *Cortinarius* was split as follows: the old genus *Cortinarius* was elevated to family level, *Cortinariaceae*, and the family was split into ten genera, of which seven were described as new. A more extensive description of the taxonomic and nomenclatural history of *Cortinarius* is provided in Supplementary Material 1.

In this paper we re-examine the dataset generated by Liimatainen et al., 2022 and argue that the genus *Cortinarius* should not yet be split due to irreproducibility of their phylogenetic analyses.

Our analyses demonstrate that the *Cortinarius* phylogeny remains unresolved and the resulting phylogenomic hypotheses suffer from very short and unsupported branches in the backbone.

This might reflect biological processes such as explosive diversification that result in the unsuitability of many or most genes to adequately reconstruct the phylogeny in an unambiguous way. While some of these problems are already visible in the original paper (poor bootstrap support for most of the branches of the backbone), further issues emerged when the data were subjected to subsequent analyses. Furthermore, we recommend that uncertainty in phylogenetic tree estimates should always be addressed and discussed in fungal phylogenomic studies, in particular when translating phylogenies to new taxonomic classifications.

## Methods

### Maximum Likelihood phylogenetic analyses

All the phylogenomic analyses are based on the concatenated fasta file, and the partition file provided by Dr. T. Niskanen (Liimatainen et al., 2022) (Liimatainen et al. 2022) including 22 sequences of 21 species (2 outgroup species, *Crepidotus sp*. BD-2015/*Hebeloma cylindrosporum* and 19 ingroup species; sequences of *C. crassus* were obtained by whole-genome sequencing and by target hybridization capture, using the same specimen), 80 loci (75 genes from (Dentinger et al. 2016) and 5 genes that are commonly used as phylogenetic markers in fungi: *RPB1, RPB2, MCM7, TEF* and *GPD*.) The outgroup species contained exclusively by 3 of the 80 loci, namely *RPB1, RPB2 and TEF1*.

Single-gene alignments were extracted from the original concatenated fasta file using RAxML-NG (v1.2.0) (Kozlov et al. 2019). We discovered eight genes where one or more species contained only one single terminal nucleotide indicating issues with the coordinates in the partition file. The single-gene alignments were therefore checked manually, and the partition file was corrected accordingly. Three supermatrices were generated for subsequent analyses: 1) original concatenation matrix: the original single-gene alignments were concatenated according to the revised partition file using PhyKIT (v1.11.7) (Steenwyk et al. 2021); 2) trimal concatenation matrix: the original single-gene alignments were trimmed using TrimAl (v1.4.1)(Capella-Gutiérrez et al. 2009) with the automated option optimized for maximum likelihood phylogenetic reconstructions and then concatenated using PhyKIT (v1.11.7); 3) gappy_50% concatenation matrix: the original alignments were trimmed using TrimAl (v1.4.1) with a gap threshold of 50% per site using the gap threshold option and then concatenated using PhyKIT (v1.11.7).

Phylogenetic analyses were performed for all datasets using the maximum likelihood (ML) criterion implemented in IQ-TREE2 (v.2.2.2.6) (Minh et al. 2020) with standard model selection by partition (Model Finder-based (Kalyaanamoorthy et al. 2017)), followed by inference (option -m TEST) using the edge-linked proportional partition model (option -p) and 10000 replicates of ultrafast bootstrap and SH-like approximate likelihood ratio test (SH-aLRT) to assess branch support. Best ML single gene trees were inferred for each dataset using IQ-TREE2 using extended model selection (option -MFP) and 1000 replicates of ultrafast bootstrap.

Tree inference for the trimal concatenation matrix was also performed with RAxML-NG (v.1.2.0)(Kozlov et al. 2019) under the GTRGAMMA model (4 discrete rate categories). During the ML search, the alpha parameter of the model of rate heterogeneity and the rates of the GTR model of nucleotide substitutions were optimized independently for each partition. Branch lengths were optimized jointly across all partitions. Non-parametric bootstrap analysis was performed on the trimal concatenation matrix, using RaxML-NG (v1.2.0). The MRE-based bootstopping test was applied after 100 replicates (Pattengale et al. 2010). Robinson-Foulds (RF) distances were used to measure topological distance between phylogenetic trees as implemented in RAxML-NG (v1.2.0)(Kozlov et al. 2019).

### Coalescent analyses and analyses of conflicting phylogenetic signal

To account for conflicting signals between concatenated and single-gene trees due to potential incomplete lineage sorting (ILS), a species tree was estimated under the multi-species coalescent (MSC) model implemented in Accurate Species TRee ALgorithm (wASTRAL-unweighted -ASTER v1.15.2.4) (Zhang et al. 2018; Zhang and Mirarab 2022) using 80 best ML likelihood single gene trees. ASTRAL is a quartet-based method and as general principle it relies on converting gene trees into unrooted four-taxon trees (called quartets) to find the summary topology that shares the maximum number of quartets with the input gene trees. Notably, branch support values are given for the quartets (quadripartitions) and not for the bipartitions (Zhang and Mirarab 2022). Average local posterior probability (LPP) (Sayyari and Mirarab 2016) on internal branches was used as branch support measure and calculated using the parameter ‘-u = 1’. LPP represents the probability that a branch is the true branch given a set of gene trees and it is a function of the number of gene trees analysed and the quartet frequencies of a branch in the species tree. Values equal or higher than 0.95 describe high confidence in a certain branch, and LPP equal to 0.7 can be considered the minimum confidence threshold for a certain branch (Sayyari and Mirarab 2016).

Given the conflicting topologies generated between methods, and particularly the uncertainty regarding the placement of the basal clade, we performed a thorough exploration of the tree space by generating 400 best-ML trees starting from 100 parsimony and 100 random starting trees in 2 independent runs using RaxML-NG and the trimal concatenation matrix. Best ML-trees were compared based on Log-likelihood score (LH) and relative RF distance between them. DensiTree (Bouckaert 2010) was used to plot the 400 tree topologies and visualize the top 4 consensus topologies. Differences between groups (run1 vs. run2 and parsimony-based vs. random starting trees) were statistically tested using a two-sided unpaired non-parametric Mann–Whitney–Wilcoxon test with a false discovery correction according to the Benjamini–Hochberg approach.

To estimate conflict among single gene trees and single gene alignment sites the gene concordance factor (gCF) (Minh et al. 2020) and the site concordance factor (sCF) (Mo et al. 2023) were estimated at each node of the best ML phylogeny generated from the trimal concatenated supermatrix using the updated maximum likelihood-based method (--scfl option) for site concordance factors implemented in IQ-TREE2 (v.2.2.2.6).

## Results and Discussion

### Re-evaluation of the phylogenomic dataset from Liimatainen et al., 2022

Based on the original concatenation matrix and partition file kindly provided by Dr. T. Niskanen, we first checked the data that should, in the ideal case, have consisted of 19 species and putatively 75 single-copy nuclear orthologs. The orthology and single-copy status was not checked in Liimatainen et al., 2022 and hence it is assumed. Five loci, *RPB1, RPB2, MCM7, TEF1* and *GPD*, commonly used fungal phylogenetic markers (Tekpinar and Kalmer 2019), were also included in the dataset but were not part of their phylogenomic analysis. However, three of them, *RPB1, RPB2 and TEF1*, were only used to root the phylogenomic tree using *Crepidotus sp*. BD-2015 and *Hebeloma cylindrosporum* as outgroup species.

Coverage was not complete for the ingroup. Short-read shallow whole-genome sequencing (WGS) yielded on average 52 genes (range 24-80 genes), whereas target hybridization capture yielded on average 68 genes (range 30-79), a difference that was just not significant (two-sided t-test with unequal variances; p = 0.08). However, the lower coverage for WGS compared to target hybridization capture highlights the potential of such technique to generate genome-scale data for fungal systematics. Excluding the outgroup species, each gene was represented by 15.2 ingroup taxa (range 6-20) and each ingroup taxon was on average represented by 61 genes (= 76%, minimum 24 genes for *C. crassus* - WGS maximum 80 genes for *C. typicus* - WGS) (Figure S1).

We combined all the available sequences into a concatenation matrix with 22 species and 80 loci and generated 3 different datasets: 1) a concatenation matrix including the original alignments with 241315 sites; 2) a concatenation matrix where the alignments were trimmed prior to concatenation using a heuristic method optimized for ML phylogenetic reconstruction implemented in TrimAl (92.44% sites of the original matrix) (Capella-Gutiérrez et al. 2009); 3) a concatenation matrix where sites with more than 50% gaps were excluded prior to concatenation (93.22% sites of the original matrix).

### Topological instability and phylogenetic conflict in *Cortinarius*

Maximum likelihood analyses for the three datasets were conducted on the partitioned data matrix using IQ-TREE2 (Minh et al. 2020) with the best-fitting model of nucleotide evolution for each partition. The resulting best ML trees were topological identical (normalized Robinson-Foulds (RF) distance = 0.0 for all pairwise comparisons) (Figure S2). For this reason, the concatenation matrix based on TrimAl alignments was used for all subsequent analyses, hereafter referred to as concatenation matrix. A best ML phylogenetic tree was also computed with RAxML-NG (Kozlov et al. 2019) but showed no topological differences with the IQ-TREE2 tree (RF=0.0) (Figure S3 A-B). To uncover potential biases introduced by inference methods or evolutionary models, a summary coalescent tree was built using 80 unrooted best ML single gene trees. Interestingly, the ML trees and the coalescent tree differed in 6 non-trivial splits (RF=0.16). Previous studies reported that species tree topological error under the MSC model can depend on the number of genes (e.g. less than 50 genes) and level of gene tree error (Sayyari and Mirarab 2016; Shekhar et al. 2018), a matter that needs further investigation for the Liimatainen et al., 2022 phylogenomic dataset.

All phylogenetic trees obtained in our analyses converged to a different topology than the one reported by Liimatainen et al., 2022 with normalized RF distance of 0.16 (all the ML trees) and 0.21 (species coalescent tree). We used four independent metrics to assess support across the different phylogenetic analyses, including non-parametric bootstrap support (BS; RAxML-NG Figure S3-A), ultrafast bootstrap (UFboot; IQ-TREE2, Figure 1B-C, S3-B), Shimodaira–Hasegawa approximate likelihood ratio test (SH-aLRT; IQ-TREE2, Figure 1B-C, S3-B), and local posterior probability (LPP; Astral, Figure 1B, S3-C).

**Figure 1.**
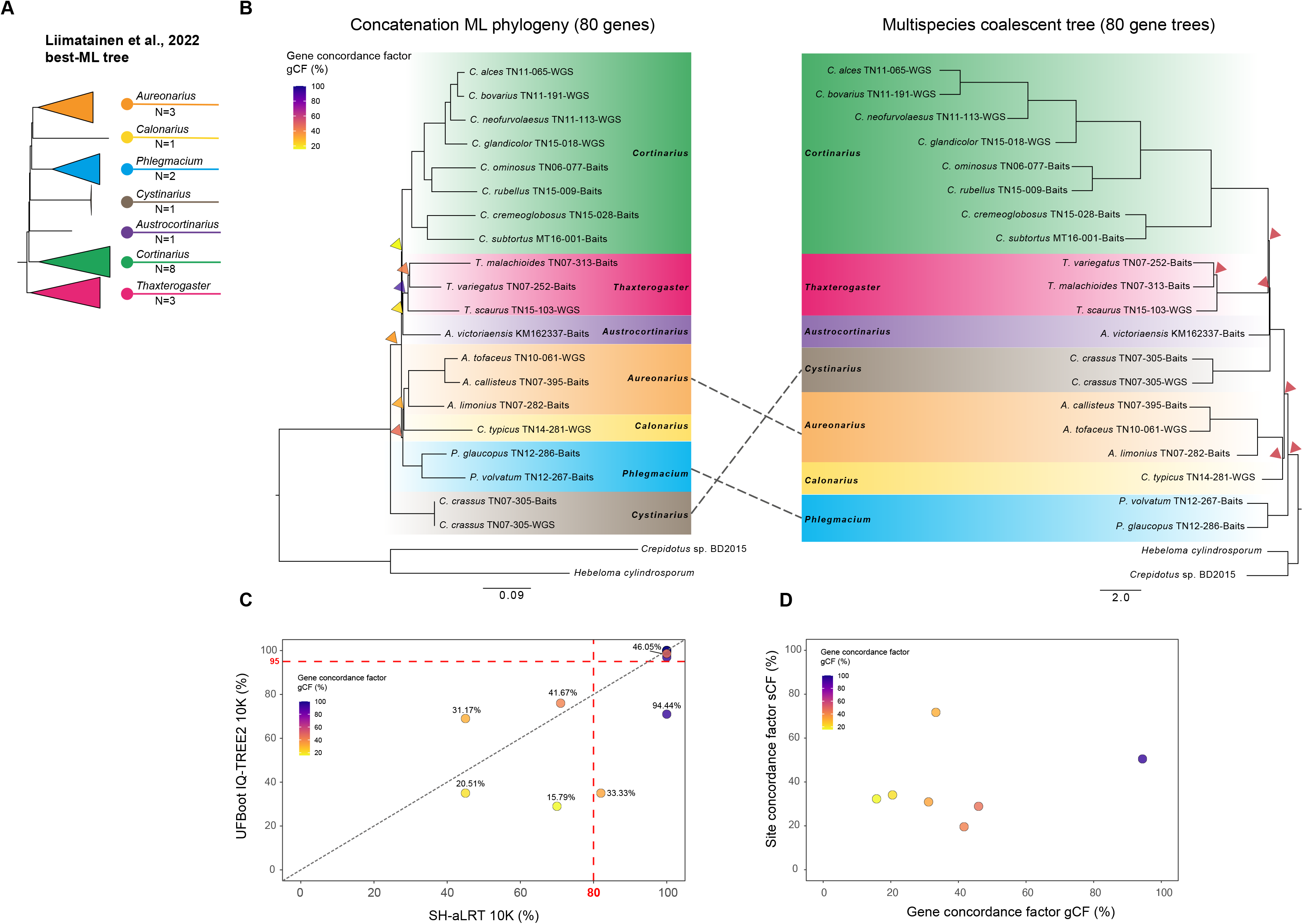
**(A)** *Cortinarius* maximum likelihood (ML) tree obtained in Liimatainen et al. (2022). The proposed new genera are collapsed according to Liimatainen et al. (2022). The number of species per genus included in the dataset are reported; **(B) Left** – Best ML tree inferred with IQ-TREE2 on the concatenation matrix of 80 putatively single-copy nuclear orthologs. Branch lengths reflect the average number of substitutions per site. **Right** – species tree generated under the multispecies coalescent model implemented in wASTRAL-unweighted using 80 unrooted best ML single gene trees. Branch lengths are reported in coalescent units. Triangles indicate 7 critical splits that received low support values using different measures: BS <80%, SH-like aLRT ≥80%, UFBoot <95% and LPP < 0.95 (see Figure S3). Triangles in the ML tree are coloured according to gene concordance factor (gCF %). Trees are rooted with *Crepidotus sp*. and *Hebeloma cylindrosporum*. **(C)** Relationship between different branch support measures: IQ-TREE2 UFBoot versus SH-like aLRT values for all branches of the ML phylogeny in (B) after 10000 generations. Dots are coloured according to gCF (%) values. The actual values are indicated next to each dot for seven critical splits. **(D)** Relationship between gCF and site concordance factor (sCF) for seven critical splits. Dots are coloured according to gCF values.

In particular, we identified seven critical splits (Figure 1 B-C-D) affecting the reliability of the backbone, showing BS <80% and either SH-like aLRT ≤80%, UFBoot ≤95% or both below the threshold. The same splits received LPP < 95% in the species coalescent tree. Out of the 7 genera, 4 are represented by more than one species and were recovered as monophyletic, *Cortinarius, Phlegmacium, Aurenoarius, Thaxterogaster*, but the branching order varies substantially across methods (Figure 1A,B,C). In the original tree BS was < 80% for *Thaxterogaster*. Six out of seven low supported splits affect the placement of the genera represented by one species only and the identity of the basal clade. Whereas all ML analyses indicate *Cystinarius* as the most basal clade (within this species selection), the species coalescent tree indicated *Phlegmacium*. Both topologies are in contrast with the phylogenomic tree reported in Liimatainen et al., 2022 (Figure 1A), where *Thaxterogaster* results as most basal group. That result was subsequently used to constrain their second tree, based on five loci (see below). Earlier phylogenetic analyses of *Cortinarius*, based mainly on ribosomal sequences (ITS and LSU regions of 16S rRNA) came to different conclusions. In the phylogeny of Peintner et al. (2004) (Peintner et al. 2004) that also had *Hebeloma* species as outgroups, the first ingroup was formed by species of *Cortinarius* in the restricted sense. The tree of Garnica et al. (2005) (Garnica et al. 2005), with *Laccaria amethystina* as outgroup, equally had *Cortinarius* in the restricted sense as first ingroup. The same result was reported by Ryberg & Matheny (2012) (Ryberg and Matheny 2012) with *Laccaria bicolor* as outgroup. A later tree by Garnica et al. (2016) (Garnica et al. 2016), though without an explicit outgroup, separated *C. crassus* (*Cystinarius*) from all remaining groups of *Cortinarius*, in agreement with our ML analyses. Soop et al. (2019)(Soop et al. 2019) on the other hand, provided two phylogenetic trees, one based on the ribosomal cluster only, and the other based on those two genes together with *RPB1* and *RPB2*. The four-gene tree, apart from several species that have not been analysed by Liimatainen et al. (2022), had species of *Thaxterogaster* as first ingroup. However, the tree based on the ribosomal genes had, apart from several species lacking in Liimatainen et al. (2022), species of *Phlegmacium* as first ingroup. Apparently, phylogenetic trees are inconsistent in resolving the relationships between the old genus *Cortinarius* in a wide sense. It is unclear to what extent these differences are caused by species selection, gene selection and/or outgroup selection.

Additionally, considering the topological differences observed between ML and coalescent trees, we estimated the degree of discordance between single-gene trees and the concatenation-based tree. We calculated the gene concordance (gCF) and site concordance factor (sCF) and investigated their relationship with UFboot and SH-aLRT (Figure 1 B,C,D). For every branch of the species tree, the gCF and sCF represent the percentage of decisive gene trees and alignment sites respectively, containing that branch (Minh et al. 2020). Discordance revolved around the six critical backbone splits identified earlier, with low gCF and sCF corresponding with splits with low BS, SH-like aLRT and UFBoot, with the exception of the of the split separating *Phlegmacium, Calonarius* and *Aurenarius* that showed SH-like aLRT=100%, UFBoot=98% but a gCF of 46% and sCF of 28.6%. Note that according to Minh et al. (2020) sCF values are seldom lower than 33%. These authors noted the particular interest of sCF values below this threshold, which may occur with high levels of incomplete lineage sorting. However, other explanations are possible and further studies into such observation are needed. This is especially pertinent considering many values are around 33% in the analysis of this data set and are associated to nodes with extremely short internal branches.

### Exploration of phylogenetic tree space

To further investigate the degree of incongruence in the dataset, we performed a thorough exploration of the trees space using RAxML-NG (Kozlov et al. 2019). In the best-case scenario, independent ML tree searches, starting from different trees, converge to a single topology with the best log-likelihood score (single likelihood peak in the likelihood space). However, depending on the dataset, distinct searches might converge to multiple topologies with large log-likelihood score differences (multiple local optima) or yield topologically highly distinct, yet almost equally likely, trees.

We inferred a total of 400 best ML trees, starting from 100 parsimony-based and 100 random starting trees, in two independent runs respectively (Figure 2). Trees were scored according to their log-likelihood. We found no significant difference in the log-likelihood distributions between runs (Wilcoxon signed rank test, *p* = 0.09), but tree searches starting from parsimony-based starting trees lead to significantly worse best ML trees than completely random ones (Wilcoxon signed rank test, *p* = 3.331e-08).

**Figure 2.**
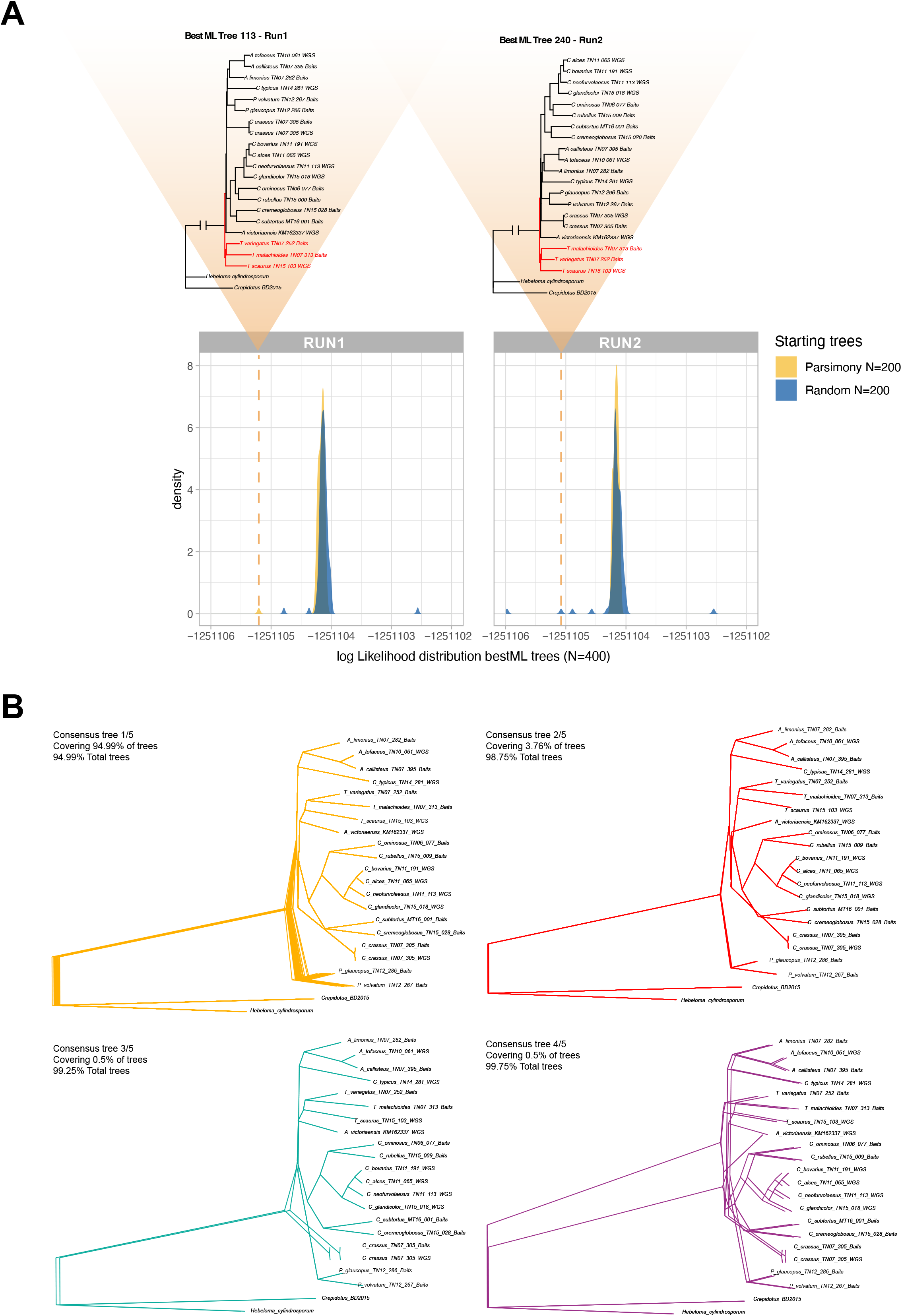
Exploration of the tree search space for the dataset generated in Liimatainen et al. (2022). **(A)** Log-likelihood score distribution for the best ML trees obtained in 2 independent runs of RAxML-NG (run 1 and run 2) starting from 200 random (blue) and 200 parsimony (yellow) starting tree per run. Triangles highlight the likelihood score of the only two topologies (one per run) that resulted in *Thaxterogaster* being the most early diverging group. **(B)** Top four consensus topologies generated in DensiTree (Bouckaert 2010) from the 400 best ML trees. The consensus 1/5 represents almost 95% of the best ML trees and corresponds to the local optima from run1 and run2 in **(A)**. Consensus 3/5 represents the 2 best ML trees (one per run) with the highest likelihood score and is topological identical to the tree reported in Figure 1B.

The two runs generated the same best ML tree that corresponds to the best ML tree generated with IQ-TREE2, showing *Cystinarius* as the most early diverging branch, although at low frequency (once in each run – that is 0.25% of the total trees). The topology generated in Liimatainen et al. (2022) with *Thaxterogaster* as basal group is only partially recovered (the genus *Austrocortinarius* branches out from *Cortinarius)* once in each run (0.25% of the total trees) (Figure 2A).

The log-likelihood distribution demonstrated the presence of a local optimum that includes 95% of best ML trees generated. ML trees within that local optimum support *Phlegmacium* as most basal clade as shown in the consensus tree generated in DensiTree (Figure 2B), in line with the multispecies coalescent tree (Figure 1B). The second consensus tree covered 3.76% of the total best ML trees and originates from the fifteen trees with the lowest log-likelihood score of the distribution. It also supports *Phlegmacium* as basal clade; however, it resolved towards *Thaxterogaster* and *Cortinarius* being descendant of *Austrocortinarius and Cystinarius*. The third and fourth consensus trees represent the two best ML trees and the two trees supporting *Thaxterogaster* as earliest diverging group respectively (Figure 2B).

Our analysis shows that the topology presented in Liimatainen et al. (2022) can be partially recovered but only as a very rare event in the tree space. Their tree is also not the best ML tree. The observation that the dataset tends to converge to one single local optimum that is also not the best ML tree merits further studies.

### Filling in the tree

Next to the 80-loci tree, Liimatainen et al. (2022) provided a second tree, based on five loci, of 245 species of *Cortinarius*. The tree was constrained with *Thaxterogaster* as outgroup, even though for both outgroup species *(Crepidotus sp. BD-2015/Hebeloma cylindrosporum)* sequences of those genes were available through previously published whole-genome sequencing. Note also that their outgroup choice was based on a tree that was not the best ML tree after concatenation (see above). Their tree added three further genera, represented by one or two species. However, not all species had sequences of all five loci. *RPB1* was represented by 243 sequences (coverage 99%) and *RPB2* by 119 sequences (coverage 49%). Coverage of the three other genes was (very) low: *MCM7* with 18/245 (7%), *GPD* with 22/245 (9%), and *TEF1* with 13/245 (5%). It is therefore safe to conclude that the 5-locus tree is largely determined by *RPB1*. At least for the three new genera only *RPB1* sequences were available.

Comparisons with earlier phylogenetic trees is only possible for the study by Soop et al. (2019) (Soop et al. 2019). However, that study contained 161 *RPB1* sequences (in a tree based on 460 sequences; 35%) and 87 *RPB2* sequences (19%). The tree signal may for that reason bear a strong signature of the ribosomal cistron, precluding a fair comparison. It may be noteworthy, though, that two of the three new genera in Liimatainen et al. (2022) are basal in the tree by Soop et al. (2019). It seems plausible that further new, probably species-poor genera will have to be added as soon as one decides to split the large genus *Cortinarius*.

### Nomenclature

The introduction of new genera did inevitably necessitate the introduction of a major number of new combinations. In fact, slightly more than half of the manuscript (43 out of 82 pages) is devoted to these new combinations on specific and intraspecific level. A few critical comments are in order.

First, their new genus *Calonarius* is threatened by *Cereicium Locquin*. Even though there is no sequence of the type species, *C. cereifolius*, the species is generally considered to be related to *C. elegantior*. Locquin (1977) (Locquin 1977) described six new genera that he segregated from *Cortinarius*, and three of them are included in Liimatainen et al. as synonyms. Of the other genera, *Hygromyxacium* (type *C. liquidus*) is likely synonym of *Cortinarius, Squamaphlegma* (type *C. aurasicaus*) likely a synonym of *Phlegmacium*, and *Cereicium* an older name for *Calonarius*.

Second, while the new combinations are all valid and legitimate according to the code of nomenclature, we cannot escape the impression that quite a few combinations were made too hastily.

That pertains to cases where earlier studies had indicated that different species, based on different types, had identical barcodes in Liimatainen et al. (2014)(Liimatainen et al. 2014). Such combinations are in our view in contravention the preamble of the code, paragraph 12 where it is stated “The only proper reasons for changing a name are either a more profound knowledge of the facts resulting from adequate taxonomic study or the necessity of giving up a nomenclature that is contrary to the rules.” The practice also deviates from Preamble point 1 that deals with the useless creation of names. In the absence of new information about the species since the publication of the barcode of the type, such names are best described as being taxonomically superfluous. There are also instances of a comparable problem, in case of names where there are no type sequences but where taxonomic practices have always considered names to be synonyms. In Liimatainen et al. (2014) such cases are species names sensu auct., a practice that differs from recommendation 50D of the Code. In Supplementary material 2 we provide an overview of the cases where this issue of taxonomic superfluity pertains.

### Generification by inflation

New insights provided by cladistic analysis can lead to a new classification that is strictly in agreement with the best estimate of the phylogeny and thereby overturns existing classifications. Such a practice demands the recognition of new genera as the previously circumscribed genus was shown to be paraphyletic or polyphyletic. Earlier subdivisions of *Cortinarius* (the traditional subgenera already recognised by Fries (1838)) were shown to be paraphyletic by Peintner et al. (2004)(Peintner et al. 2004) and Garnica et al. (2005)(Garnica et al. 2005), and it was therefore inevitable that these genera had to be given up. In the study of Liimatainen et al. (2022) the authors purported to recognise monophyletic groups within the monophyletic *Cortinarius* as previously described. The authors could therefore have chosen to recognise their monophyletic groups below the level of genus (and hence keeping the monophyletic *Cortinarius*) or to recognise these groups on generic rank (and then elevating the genus *Cortinarius* to family level). This practice has been referred to by Kuyper (1994) (Kuyper 1994) as generification by inflation. How necessary is such a practice in the case of *Cortinarius*?

We agree with Liimatainen et al. (2022) that ranking is to a certain extent a subjective process and that mycologists have to strike a balance between the number of genera recognised and the amount of diversity they include. The authors noted a general tendency of making smaller genera as a result of molecular phylogenetic analysis. Of the three cases they mention in that respect, two refer to previous classifications that had to be given up because they contained paraphyletic or polyphyletic groups, and one where a classification into one family with seven genera is exactly equivalent to a classification of one genus with seven subgenera. In that case one could equally ask the question to what extent this constitutes generification by inflation.

This ghost of taxonomic inflation is far from imaginary(Isaac et al. 2004; Sandall et al. 2023). Liimatainen et al. (2022) provided reflections on why they include the /carbonellus clade in *Phlegmacium*, as otherwise the latter genus would have to be split in four genera. They also showed that one could easily split *Cortinarius* in their restricted sense, as it still is by far the largest of their genera with likely over 70% of currently known species, into eleven genera. Combined with a poorly resolved phylogeny without support for the backbone, and the occurrence of several species that have an isolated position in the currently published phylogenies, it is difficult to see how this splitting process can be stopped easily as some splits will likely generate paraphyletic groups. One would wonder, for instance, what would happen with subgenus *Orellani*, where the two species, *C. orellanoides* and *C. rubellus* do not form a monophyletic group, even though both names are generally considered to be synonyms. And with an ever-growing number of future genera in this clade we would again be confronted with the lack of hierarchical structure in such a classification.

Vellinga et al. (2015) (Vellinga et al. 2015) listed five guidelines and one recommendation (for editors of scientific journals) when introducing new genera. These guidelines include the need to have all (new) genera monophyletic and the evidence for the monophyly of these groups should be based on more than one gene. It is evident that the paper by Liimatainen et al. (2022) fully complies with those guidelines. It also seems that the authors of Vellinga et al. (2015) were too optimistic about the benefits of including more genes. Quantifying the distribution of phylogenetic signal in phylogenomic datasets it has become as important as generating the phylogenomic datasets itself, considering that a handful of genes can sometimes drive the resolution of contentious nodes (Shen et al. 2017).

A third guideline related to the species to be included in the tree, such as inclusion of type species, sufficient geographical coverage and a representative species selection. Here the paper by Liimatainen et al. (2022) does not fare well. Most type species have not been included, geographical coverage, especially of the 80-loci tree is limited, with 18 (of 19 species) studied occurring in the northern hemisphere, and several species that are basal in the phylogeny by Soop et al. (2019) equally lacking in the 80-loci tree.

The next guideline would be to only translate a phylogeny into a classification if the branches have credible statistical support. In this case the backbone of the tree remains unresolved and different methods (concatenation versus coalescence; tree searching programs) yielded different basal groupings. Vellinga et al. (2015) also stated that a list of options should be given, and alternatives be tested. The only reference in Liimatainen et al. (2022) to alternatives is about alternative forms of splitting the genus. The only arguments they provide for not splitting the genus is nomenclatural stability and the fact that the genus *Inocybe* has been split, a split that could equally fall under the category of taxonomic inflation. We in contrast argue that the very poor resolution of the phylogeny caused by extansive gene-tree/species-tree confilicts and a rather unrepresentative species selection clearly supports the famous dictum “in dubiis abstine”, in case of doubt refrain from major taxonomic and nomenclatural changes.

### If phylogenomics cannot (yet) resolve the *Cortinarius* phylogeny, what’s next?

Our new analysis has shown even more clearly that the backbone of the *Cortinarius* phylogeny is very poorly supported and consequently the relationships between the clades cannot (easily) be resolved. The explanation for our inability to provide a credible solution to the phylogeny of *Cortinarius* is to be found in the very short backbone branches in the genus. These short branches were already noted by Peintner et al (2004) (Peintner et al. 2004) but in that study attributed to low DNA divergence. The analysis of the 80-loci tree with the (frequently very) low gene concordance and site concordance factors would rather suggest the opposite explanation. That alternative explanation fits with data by Ryberg & Matheny (2012) (Ryberg and Matheny 2012) who reported much higher nucleotide substitution rates in *Cortinarius* compared with almost all other groups of ectomycorrhizal Agaricales. The very low sCF factors, often close to 33%, the theoretical minimum, in the basal branches of the tree is equally an indicator for considerable randomness (or lack of phylogenetic structure) in the data. One possible solution would then be to be more selective in choosing genes for phylogenomic analysis. We could then try to select those genes based on sequence-, tree-, and function-based properties (Shen et al. 2016a). However, the current data could not be applied for that calculation as we do not have sequence data for all loci for all species. There is a need for additional data before this analysis can be executed.

However, one should also be open to the alternative possibility, viz., that the inability of providing a clear resolution of the *Cortinarius* tree is inherent in the data, a problem that then could not be resolved with more species, or more (or: less and better) genes. Short backbone branches are likely a reflection of rapid diversification or explosive speciation after the genus evolved. Such rapid diversification has often been interpreted as adaptive radiation, but Givnish et al., 2015 (Givnish 2015) indicated the need for a conceptual distinction. Adaptive radiation is caused by competition between closely related and ecologically similar species resulting in divergent selection and the evolution of key innovations that allows more efficient utilisation of niches. One might consider the evolution of the ectomycorrhizal habit in *Cortinarius* as a key innovation, as has been suggested for several groups of ectomycorrhizal fungi (Sánchez-García and Matheny 2017; Sánchez-García et al. 2020; Sato 2024). However, the process of niche filling would ultimately reduce speciation rates, but this has not been found, and in some groups of *Cortinarius* even increasing speciation rates have been noted (Ryberg and Matheny 2012).

While we currently do not know the factors that caused the high diversification rates (and nucleotide substitution) rates in *Cortinarius*, a consequence of rapid diversification is likely an increased chance of incomplete lineage sorting. Incomplete lineage sorting is likely a major explanation for these very short basal branches in the backbone phylogeny. Hybridization, although only infrequently reported for Agaricales, could be a further cause of very short branches. Under such conditions of short branches due to rapid speciation events concatenation-based methods will have a high likelihood of leading us to the incorrect tree (Whitfield and Lockhart 2007; Schrempf and Szöllősi 2020).

## Conclusions

With high likelihoods of producing incorrect trees, we should be careful in prematurely proposing radical taxonomic and nomenclatural changes. For that reason, we argue that the genus *Cortinarius* should not (yet) be split.

## List of abbreviations

MSC: multispecies coalescent
rDNA: ribosomal DNA
ML: maximum likelihood
SH-aLRT: SH-like approximate likelihood ratio test
RF: Robinson-Foulds
ILS: incomplete lineage sorting
LPP: local posterior probability
gCF: gene concordance factor
sCF: site concordance factor
BS: bootstrap
UFBoot: Ultra-fast bootstrap

## Ethics approval and consent to participate

Not applicable

## Adherence to national and international regulations

Not applicable

## Consent for publication

All three authors have seen and fully agreed with the current submitted version. The manuscript has not been submitted, in parts or as a whole, to other journals; nor do we intend to submit (parts of) the manuscript to other journals as long as the manuscript is under consideration by IMA fungus.

## Availability of data and materials

Alignments and phylogenetic trees generated in the current study are available at the following Dryad repository: DOI: 10.5061/dryad.m63xsj492 (in progress)

## Competing interests

Not applicable

## Funding

Marie-Skłodowska Curie Individual Postdoctoral Fellowship program (no. HORIZON-MSCA-2021-PF-01, 101065406-GLiMMer) to Brigida Gallone

## Authors’ contributions

All three authors have conceptualised the study, contributed to the analysis and interpretation of the various data, and have written parts of the original drafts.

## Acknowledgements

We thank the European Commission for providing funding to Brigida Gallone under the Marie-Skłodowska Curie Individual Postdoctoral Fellowship program (no. HORIZON-MSCA-2021-PF-01, 101065406-GLiMMer).

---

**Figure S1.** Occupancy matrix showing presence/absence of genes (x-axis) per species (y-axis) in the supermatrix generated in Liimatainen et al. (2022). Species are grouped according to Figure 1.

**Figure S2.** Best maximum likelihood trees inferred with IQ-TREE2 on the concatenation matrix of 80 putative single-copy nuclear orthologs used in Liimatainen et al. (2022). The single gene alignments were differentially trimmed before concatenation and tree inference: **(A)** original concatenation matrix: the original single-gene alignments were concatenated according to a revised partition file; **(B)** gappy 50% concatenation matrix: the original alignments were trimmed using TrimAl (v1.4.1) with a gap threshold of 50% per site using the gap threshold option; **(C)** trimal concatenation matrix: the original single-gene alignments were trimmed using TrimAl (v1.4.1) with the automated option optimized for maximum likelihood phylogenetic reconstruction; SH-like aLRT and UFBoot support values for 10000 generations are indicate for each branch. Trees are rooted with *Crepidotus sp*. and *Hebeloma cylindrosporum*. Branch lengths reflect the average number of substitutions per site. Genera are colored and grouped based on in Liimatainen et al. (2022).

**Figure S3. (A)** Best ML tree inferred with RAxML-NG on the concatenation matrix of 80 putative single-copy nuclear orthologs. Number on branches indicate non-parametric bootstrap for 100 generations. Branch lengths reflect the average number of substitutions per site. **(B)** Best ML tree inferred with IQ-TREE2 on the concatenation matrix of 80 putative single-copy nuclear orthologs. Branch lengths reflect the average number of substitutions per site. Number on branches indicate SH-aLRT support (%) / ultrafast bootstrap support (%)/gCF (%) /sCF (%). The tree is the same as Figure 1B but it includes full annotations of the support values. **(C)** Species tree generated under the multispecies coalescent model implemented in wASTRAL-unweighted using 80 unrooted best ML single gene trees. Branch lengths are reported in coalescent units. Number on branches indicate local posterior probabilities (LPP, decimals from 0.01 to 0.99). Triangles indicate 7 critical splits with LPP < 95%. The tree is the same as Fig1B but it includes full annotations of the support value.

## Notes

### Competing Interest Statement

The authors have declared no competing interest.

