## Supplemental Note 1 for "The genus *Cortinarius* should not (yet) be split"

**Supplemental Material 1**

### A short history of *Cortinarius*

As a genus *Cortinarius* is generally easily recognisable by the combinations of a rusty brown spore print, ornamented spores and the presence of a cortina, a cobweb-like structure that connect the pileus with the stipe in young specimens. Fries (1851) already considered the genus as a natural (in the pre-Darwinian sense of common essence, not of common descent) genus. The description of *Cortinarius* by Fries (1838) still fits the overwhelming majority of species that are currently classified in *Cortinarius*. Fries (1821) mentioned four tribus under his series *Cortinaria*, viz., *Telamonia*, *Inoloma*, *Phlegmacium* and *Dermocybe*. He added a fifth tribus, *Myxacium*, under series *Derminus*. Fries (1838) subdivided his genus, in which he then placed 216 species, in six subgroups that have usually been considered subgenera, but sometimes genera. These infrageneric taxa were the four taxa earlier listed under series *Cortinaria*, to which he added *Myxacium* and a new infrageneric taxon. The six infrageneric taxa are *Myxacium*, *Phlegmacium*, *Inoloma*, *Dermocybe*, *Telamonia* and *Hydrocybe*. In most cases these infrageneric groups are easily recognisable and therefore still appear as separate groups in identification keys. Only separating *Telamonia* and *Hydrocybe* was considered too complex and they have more often been merged. This classification was in fact eurocentric and did not undergo major modifications. Only *Sericeocybe* (a segregate from *Inoloma*) and *Leprocybe* (a segregate from *Dermocybe*) have been added. This subdivision of *Cortinarius* in fairly easily recognisable subgenera worked well for the *Cortinarius* species of the northern hemisphere, and especially Europe. It still forms the basis for almost all practical identification keys for northern-hemisphere species of *Cortinarius*. Integrating species from the southern hemisphere turned out to be more challenging. In a study of *Cortinarius* species in the southern part of South America (the area dominated by trees of the Nothofagaceae), Moser & Horak (1975) added two further subgenera (*Paramyxacium* and *Cystogenes*), but refrained from a more radical modification of an infrageneric classification. They also noted linkages between *Cortinarius* and several other genera such as the sequestrate genus *Thaxterogaster*, and the genera *Rozites* and the new genus *Stephanopus*. Under the then current paradigm that agaricoid fungi were derived from gastroid and sequestrate forms, Moser & Horak (1975) noted that this paradigm would necessitate the independent evolution of *Cortinarius*, as conceived by them. They stated explicitly that such an evolutionary scenario seemed extremely unlikely to them and therefore proposed the alternative scenario that such gastroid and sequestrate forms were derived from *Cortinarius*. The implication would be that *Thaxterogaster* would be an unnatural grouping within *Cortinarius*, but at that time they refrained from drawing the nomenclatural consequences.

The development of methods to generate DNA sequences and apply those sequences in an evolutionary context allowed attempts to revisit the existing *Cortinarius* classification. Initial studies by Liu et al. (1997) Høiland and Holst-Jensen (2000), and Moncalvo et al. (2002), based on ITS and / or LSU sequences, confirmed that *Cortinarius* was monophyletic. More importantly these studies established that many of the traditionally recognised subgenera (*Myxacium*, *Phlegmacium*, *Telamonia* inclusive of *Hydrocybe*) were polyphyletic. No attempt was made in these studies to propose a new classification based on monophyletic subgroups that could then be recognised on subgeneric or generic level. On the other hand these studies contributed to the enlargement of the genus *Cortinarius*. Peintner et al. (2001) intended to establish relations between 'true' *Cortinarius* species and secotioid and gastroid mushrooms that were considered by Moser & Horak (1975) to have been derived from *Cortinarius*. Their study, based on the analysis of 151 ITS sequences, indicated multiple origins of these sequestrate and gasteroid forms that were all nested within *Cortinarius*. For that reason the genera *Thaxterogaster*, *Protoglossum* and *Quadrispora* were subsumed under *Cortinarius*. The study also showed that both *Thaxterogaster* and *Protoglossum* were polyphyletic, a conclusion already anticipated by Moser & Horak (1975). A further study (Peintner et al., 2002), also based on ITS sequences, indicated that the genera *Rozites*, *Cuphocybe* and *Rapacea*, which were considered to be genera closely related to *Cortinarius*, were equally nested in *Cortinarius*. That study also showed that both *Rozites* and *Cuphocybe* were polyphyletic. The genus *Stephanopus* was not represented by any sequences (a situation that has not changed since) and its relation with *Cortinarius* remains unknown.

A first attempt to arrive at a phylogenetically supported new classification of *Cortinarius* was undertaken by Peintner et al. (2004). Based on 186 ITS and 54 LSU sequences they noted fourteen well-supported (Bayesian Posterior Probabilities > 70%; note that half of these clades received less than 50% bootstrap support) clades with three or more species, next to several singleton species that constituted further clades. They also stated that their molecular-sequence data demonstrated that species of *Cortinarius*, although morphologically quite variable, are conserved by comparison in their rDNA region. Short basal branches characterize all *Cortinarius* phylogenies. They interpreted these short branches as reflecting the low divergence found in the ribosomal RNA gene. We discuss the relationship between short basal branches and low divergence in the main body of the manuscript.

As a consequence of this analysis they confirmed that the extended concept of *Cortinarius* was monophyletic. They did not formally propose a new classification, and ended their paper with the question 'Can the *Cortinarius* phylogeny be resolved?'. They provided a cautious and provisional 'not yet', suggested that more extensive taxon sampling and more extensive character sampling (i.e., more than two genes of the ribosomal cluster) are needed for a better resolved phylogeny that could be translated in a new classification.

Subsequent efforts worked on increasing taxon sampling and / or character sampling. Garnica et al. (2005) extended the study to 262 species, with a somewhat larger representation of species from the southern hemisphere. They only used two genes, viz., ITS and LSU, sequenced in one stretch. Whereas their tree with eight major clades shows similarities with the earlier classification by Peintner et al. (2004), it also shows some notable differences, especially in the basal position of the /telamonia clade. It is not clear whether their outgroup choice (*Laccaria*) did impact on that tree structure. Basal branches in the tree usually received little or no statistical support. Stensrud et al. (2014) applied three genes, all from the ribosomal cluster (SSU, ITS, LSU) but focused mostly on taxa from the northern hemisphere (81 species; only five species from the southern hemisphere were included). The study recognised twelve monophyletic groups and a number of singleton species. Many of their monophyletic groups were also recovered by Peintner et al. (2004). They refrained from formally naming those clades. They finally noted that basal branches of the tree received very little support, but did not provide an explanation for that observation.
A major step forward was the analysis of Soop et al. (2019). They used a dataset of 730 species for which both ITS and LSU data were available and a second dataset with 460 species for which sequences of two additional genes (RPB1 and RPB2) were available. Their dataset also showed a much better representation of *Cortinarius* species from the southern hemisphere, especially from Australasia; South America and especially Africa were almost completely absent from their species listing. Soop et al. formally recognised 79 clades that they formally described as sections and recovered an additional 20 clades where they refrained from formal description. They noted that the 4-locus tree in most cases recovered the same clades as the 2-locus tree, but often increased statistical support. Especially the addition of RPB1 often enhanced clade support. No further taxonomic hierarchy was proposed. They stated that for a more complete hierarchical framework that uses these sections as building blocks for a complete hierarchical structure a larger number of genes or preferably a phylogenomics approach would be required.
That step was recently taken by Liimatainen et al. (2022). They reported to have performed phylogenomic analysis for 19 species based on 75 single-copy genes; and a subsequent, constrained analysis for 245 species based on five single-copy genes (see the main body of the paper for the actual numbers). On the basis of that analysis they decided to split the genus *Cortinarius*. The old genus *Cortinarius* was elevated to the family level, Cortinariaceae. The family was split into ten genera, of which seven were described as new genera. Four new genera were erected based on whole-genome sequencing and baiting, and three further genera were proposed as a result of the analysis of five single-copy genes. They also introduced several new subgenera and sections and, as a consequence of their splitting the old genus *Cortinarius*, a very considerable number of section (41) and species (541) names.

The strength of their paper is that they used a combination of (admittedly shallow) whole-genome sequencing and the use of baits to capture specific genes. Especially that latter method holds great promise in generating sufficient data for multigene analyses.
