## Supplemental Note 2 for "The genus *Cortinarius* should not (yet) be split"

**Supplementary Material 2**

**Nomenclatural superfluities**

Here we mention cases of new combinations for taxa where previous ITS barcoding (often by authors that are part of the current paper) have indicated identical barcodes of the type collection. We refer to these novelties as taxonomically superfluous combinations, a term that has no standing under the rules of nomenclature, but refers to combinations that are in contravention of the preamble of the code, paragraph 12 where it is stated "*The only proper reasons for changing a name are either a more profound knowledge of the facts resulting from adequate taxonomic study or the necessity of giving up a nomenclature that is contrary to the rules*." The practice also deviates from Preamble point 1 that deals with the useless creation of names. In the absence of new information about the species since the publication of the barcode of the type, such names are best described as being taxonomically superfluous. There are also instances of a comparable problem, in case of names where there are no type sequences but where taxonomic practices have always considered names to be synonyms. In Liimatainen et al. (2014) (Liimatainen et al. 2014) such cases are species names sensu auct., a practice that differs from recommendation 50D of the Code. These two categories are listed separately.

We refer to the species names under *Cortinarius*, as we do not think there is currently sufficient evidence for splitting the genus.

*Cortinarius acidophilus* - a synonym of *C. pseudonaevosus* (recombined in a subsequent paper)

*Cortinarius barbarorum* - a synonym of C. *metarius* (also recombined in their paper)

*Cortinarius brunneoviolaceus* - a synonym of *C. brunneolividus* (also recombined in their paper)

*Cortinarius cephalixolargus* - a synonym of *C. largus* (combination already existing)

*Cortinarius cinctipes* - a synonym of *C. cliduchus* (also recombined in their paper)

*Cortinarius clarus* - a synonym of *C. largus* (combination already existing)

*Cortinarius concrescens* - a synonym of *C. balteatoalbus* (combination already existing)

*Cortinarius congeminus* - a synonym of *C. largus* (combination already existing)

*Cortinarius corrugis* – a synonym of *C. turmalis* (also recombined in their paper)

*Cortinarius crenulatus* - a synonym of *C. talus* (also recombined in their paper)

*Cortinarius cupreoviolaceus* - a synonym of *C. largus* (combination already existing)

*Cortinarius eumarginatus* - a synonym of *C. purpurascens* (also recombined in their paper)

*Cortinarius frondosophilus* - a synonym of *C. platypus* (also recombined in their paper)

*Cortinarius genuinus* - a synonym of *C. collocandoides* (also recombined in their paper)
*Cortinarius josephii* - a synonym of *C. gracilior* (combination already existing)
*Cortinarius largoides* – a synonym of *C. subpurpurascens* (also recombined in their paper)
*Cortinarius luteovaginans* - a synonym of *C. aurantiopallidus* (also recombined in their paper)
*Cortinarius misermontii* – a synonym of *C. olidoamarus* (also recombined in their paper)
*Cortinarius muricinicolor* - a synonym of *C. variecolor* (combination already existing)
*Cortinarius mutabilis* - a synonym of *C. occidentalis* (also recombined in their paper)
*Cortinarius ochropudorinus* - a synonym of *C. talus* (also recombined in their paper)
*Cortinarius olidus* – a synonym of *C. cliduchus* (also recombined in their paper)
*Cortinarius parolivascens* - a synonym of *C. scaurus* (also recombined in their paper)
*Cortinarius piriodolens* - a synonym of *C. variecolor* (combination already existing)
*Cortinarius pseudocephalixus* – a synonym of *C. cliduchus* (also recombined in their paper)
*Cortinarius pseudominor* - a synonym of *C. talus* (also recombined in their paper)
*Cortinarius pseudopansa* - a synonym of *C. varius* (combination already existing)
*Cortinarius pseudopimus* - a synonym of *C. varius* (combination already existing)
*Cortinarius pseudotalus* - a synonym of *C. talus* (also recombined in their paper)
*Cortinarius pseudoturmalis* - a synonym of *C. claricolor* (combination already existing)
*Cortinarius rufior* - a synonym of *C. varius* (combination already existing)
*Cortinarius saginoides* - a synonym of *C. varius* (combination already existing)
*Cortinarius subaccedens* – a synonym of *C. olidoamarus* (also recombined in their paper)
*Cortinarius subamaricatus* - a synonym of *C. tirolianus* (also recombined in their paper)
*Cortinarius subcyanites* - a synonym of *C. cyanites* (combination already existing)
*Cortinarius subdecoloratus* - a synonym of *C. ochraceobrunneus* (also recombined in their paper)
*Cortinarius subfuligineus* - a synonym of *C. subrugulosus* (also recombined in their paper)
*Cortinarius subinops* - a synonym of *C. subpurpurascens* (also recombined in their paper)
*Cortinarius subvariiformis* - a synonym of *C. luteocingulatus* (also recombined in their paper)
*Cortinarius thalliopurpurascens* - a synonym of *C. herpeticus* (also recombined in their paper)
*Cortinarius vacciniophilus* - a synonym of *C. pseudonaevosus* (but only recombined in a subsequent paper)

The second category pertains to names that are likely potentially taxonomically superfluous. These cases involve species names that have not been typified but have been and still are in current use, and where the taxonomic interpretation does not seem to be in doubt. Such cases have been indicated in Liimatainen et al. (2014) (Liimatainen et al. 2014) as sensu auct. This terminology is potentially misleading, as it involves a different interpretation of that concept than specified in the rules of nomenclature (Recommendation 50D, where misidentifications are referred to as misapplications, which should be indicated by “auct., non [followed by the name of the original author]. In some cases Liimatainen et al. (2022) (Liimatainen et al. 2022) accepted the names in its current interpretation and made new combinations (e.g., *C. vespertinus*, where the possible new combination based on *C. variipes* has not been made; *C. sulfurinus*, for which no synonyms were reported), while in other cases they combined both the name in current use (s. auct.) and very likely younger synonyms. These cases are:

*Cortinarius alnobetulae* - a synonym of *C. moseri* s. auct. (also recombined in their paper)
*Cortinarius calojanthinus* - a synonym of *C. corrosus* s. auct. (also recombined in their paper)
*Cortinarius elotoides* - a synonym of *C. pseudoglaucopus* s. auct. (also recombined in their paper)
*Cortinarius evosmus* - a synonym of *C. osmophorus* s. auct. (also recombined in their paper)
*Cortinarius flavescentipes* - a synonym of *C. balteatocumatilis* s. auct. (also recombined in their paper)
*Cortinarius gentianeus* - a synonym of *C. caesiostramineus* s. auct. (combination already existing)
*Cortinarius juxtadibaphus* - a synonym of *C. dibaphus* s. auct. (also recombined in their paper)
*Cortinarius latoclaricolor* - a synonym of *C. durus* s. auct. (also recombined in their paper)
*Cortinarius leonicolor* - a synonym of *C. amoenolens* s. auct. (also recombined in their paper) and of *C. anserinus* s. auct. (combination already existing)
*Cortinarius mendax* - a synonym of *C. subporphyropus* s. auct. (also recombined in their paper)
*Cortinarius neotriumphans* - a synonym of *C. saginus* s. auct. (also recombined in their paper)
*Cortinarius ophiopus* - a synonym of *C. triumphans* s. auct. (combination already existing)
*Cortinarius rapaceoides* - a synonym of *C. caroviolaceus* s. auct. (also recombined in their paper)
*Cortinarius scaurocaninus* - a synonym of *C. magicus* s. auct. (also recombined in their paper, but in *Thaxterogaster*, whereas *C. scaurocaninus* has been transferred to *Phlegmacium*)
*Cortinarius subpurpureophyllus* - a synonym of *C. napus* s. auct. (also recombined in their paper)
*Cortinarius triumphalis* - a synonym of *C. vulpinus* s. auct. (combination already existing)
*Cortinarius veneris* - a synonym of *C. balteatocumatilis* s. auct. (also recombined in their paper)

Finally, we list two cases of evident generic misclassifications. *Cortinarius coniferarum* was reclassified in *Calonarius*. Its basionym, *C. multiformis* var. *coniferarum*, is generally considered closely related to or a synonym of *C. multiformis*, as species classified in *Thaxterogaster*. *Cortinarius magicus* was reclassified in *Thaxterogaster*. The species is generally considered a synonym of *C. scaurocaninus*, classified in *Phlegmacium*.
