## Supplementary figures and images for "The genus *Cortinarius* should not (yet) be split"

### Figure S1

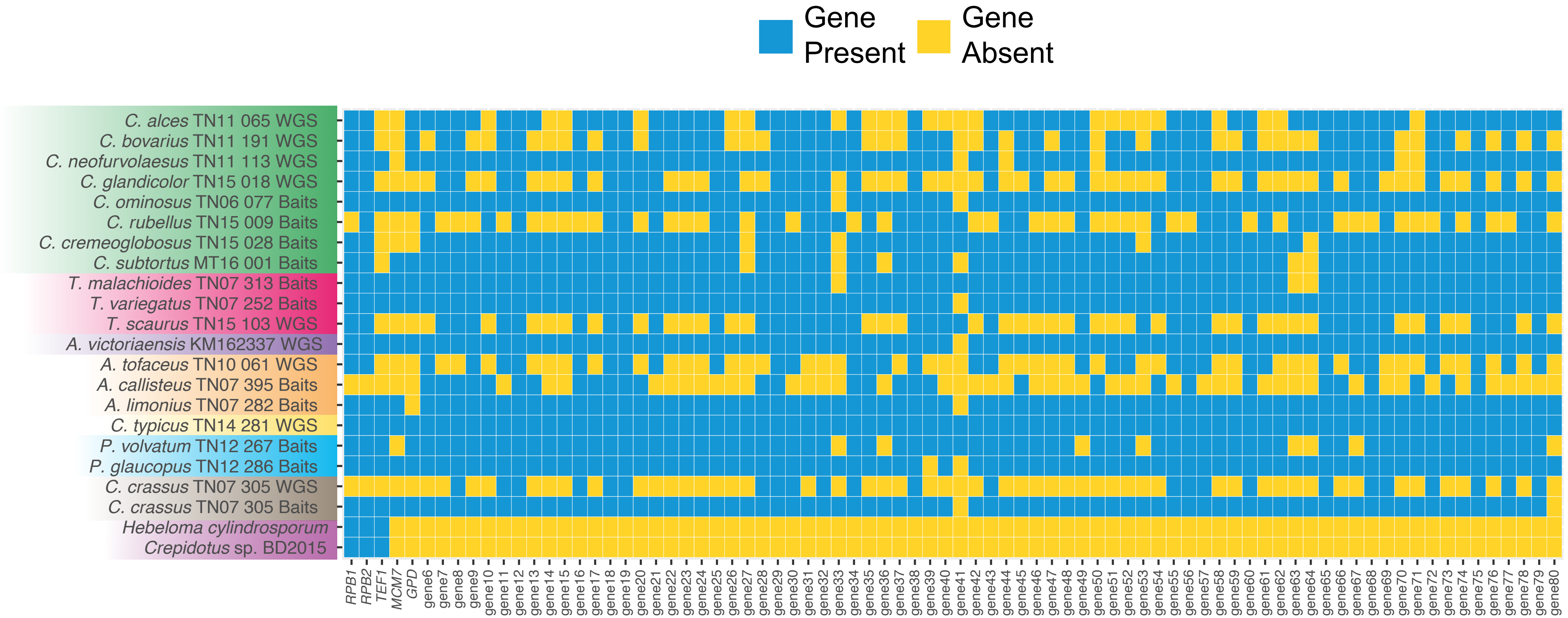

### Figure S2

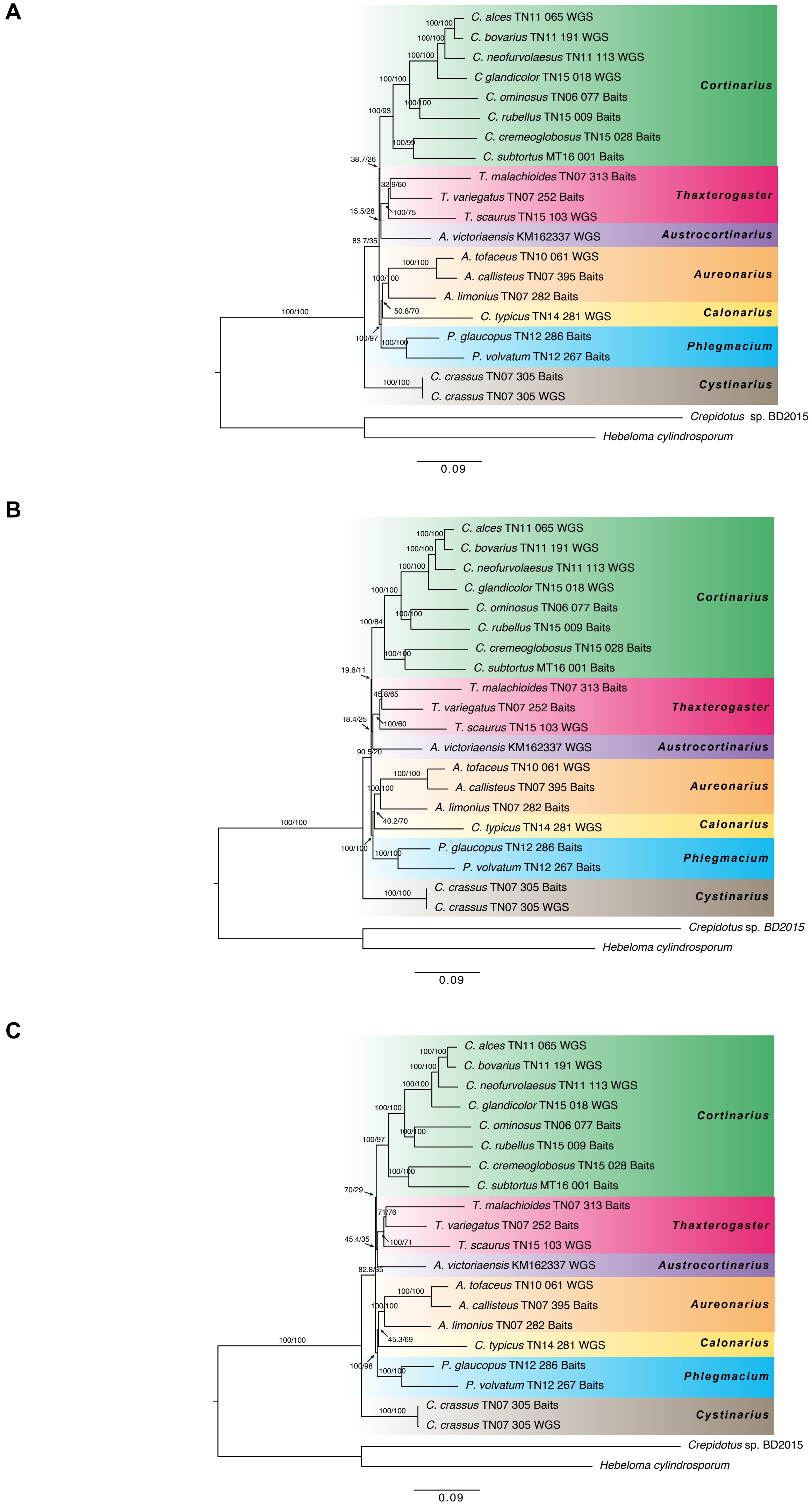

### Figure S3

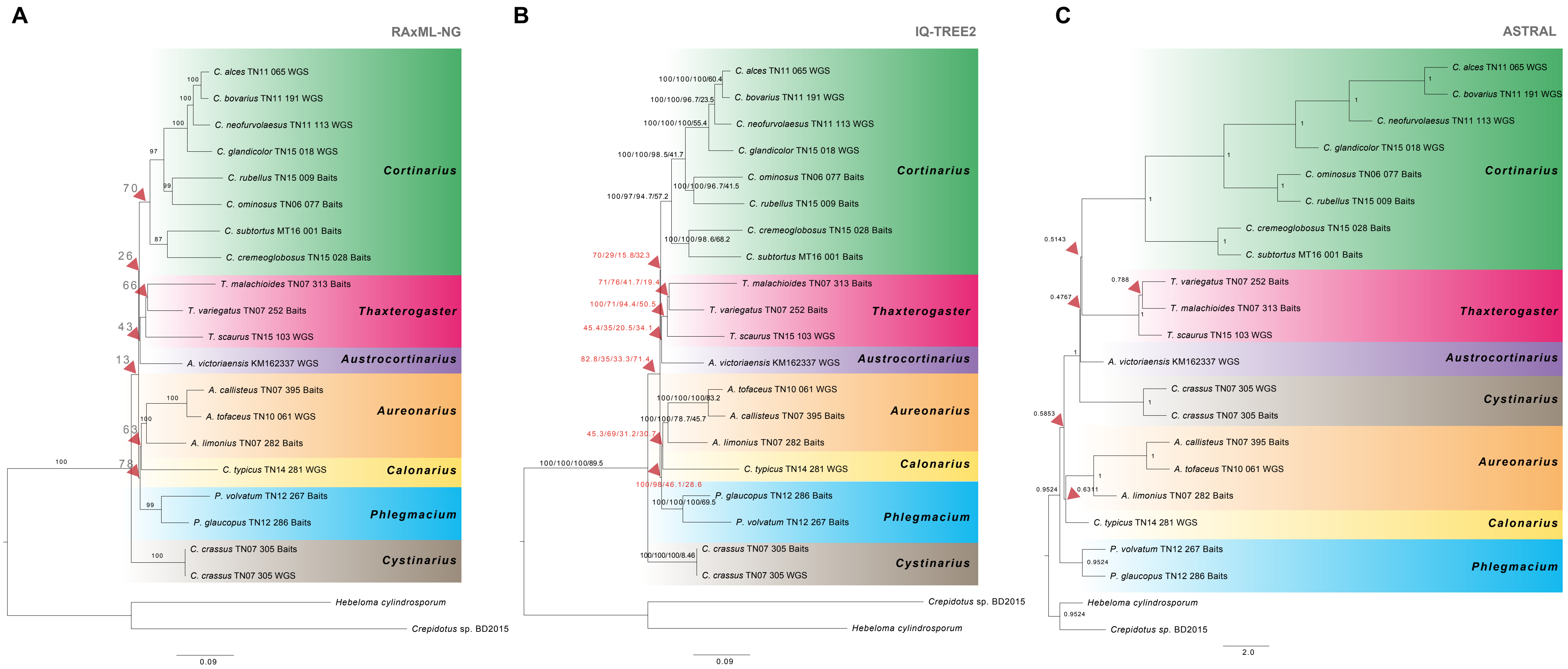
